# Tumor microenvironment noise-induced polarization: the main challenge in macrophages’ immunotherapy for cancer

**DOI:** 10.1101/2024.05.02.592270

**Authors:** Jesus Sierra, Ugo Avila-Ponce de León, Pablo Padilla-Longoria

## Abstract

Disturbance of epigenetic processes can lead to altered gene function and malignant cellular transformation. In particular, changes in the epigenetic landscape are a central topic in cancer biology. The initiation and progression of cancer are now recognized to involve both epigenetic and genetic alterations. In this paper, we study the epigenetic mechanism (related to the tumor microenvironment) responsible for increasing tumor-associated macrophages that promote the occurrence and metastasis of tumor cells, support tumor angiogenesis, inhibit T cell-mediated anti-tumor immune response, and lead to tumor progression. We show that the tumor benefits from the macrophages’ high degree of plasticity and larger epigenetic basins corresponding to phenotypes that favor cancer development through a process that we call noise-induced polarization. Moreover, we propose a mechanism to promote the appropriate epigenetic stability for immunotherapies involving macrophages, which includes p53 and APR-246 (eprenetapopt). Our results show that a combination therapy may be necessary to ensure the proper epigenetic stability of macrophages, which otherwise will contribute to cancer progression. On the other hand, we conclude that macrophages may remain in the anti-tumoral state in types of cancer that exhibit less TP53 mutation, like colorectal cancer; in these cases, macrophages’ immunotherapy may be more suitable. We finally mention the relevance of the epigenetic potential (Waddington’s landscape) as the backbone for our study, which encapsulates the biological information of the system.

## 1. Introduction

The tumor microenvironment (TME) supplies a fundamental ecological niche for cancer initiation and development. Although there is considerable diversity in the inflammatory components of the TME in cancers from different tissues, the common denominator is the infiltration of monocytes, macrophages, and dendritic cells [19]. Macrophages are particularly interesting to study from a stochastic analysis point of view due to their plasticity in response to environmental signals. The functional differences of macrophages are closely related to their plasticity. Moreover, molecules in TMEs are responsible for regulating macrophages’ functional phenotypes [25]. Such molecular signals are so diverse and random that we consider it fit to treat them as Gaussian noise that increases in magnitude as the tumor progresses. Under this assumption, our mathematical model shows that most tumor-associated macrophages (TAMs) get eventually polarized into macrophages with phenotypes that favor cancer development (these phenotypes have the largest epigenetic basins); this result agrees with the observation in [40] of adverse effects of TAMs on overall survival of patients with gastric, breast, bladder, ovarian, oral, and thyroid cancers, except for positive outcome in patients with colorectal cancer. The latter result may indicate the influence of p53 on macrophages: in [20], 40-60% of colorectal cancer tumors show TP53 mutation, whereas in ovary cancer, it is 90%. We will discuss later the relevance of p53 on immunotherapies involving macrophages.

Our model shows that the time required for noise-induced polarization of macrophages depends on the noise’s magnitude. Hence, we should have fewer macrophages with pro-tumoral features in the early stages of cancer or in situations where the noise associated with the TME is weaker; this may explain why in [32], macrophages in early-stage human lung cancer did not exert immunosuppressive functions.

## 2. Materials and methods

### 2.1. Network Reconstruction, Boolean States, and Epigenetic Landscape

We start our analysis by considering a concise network of the molecular basis of macrophage polarization and the Boolean attractors obtained from it. Both sources of data were previously reported in [2, 3]. Our reconstructed network was obtained through a bottom-up approach, where we set up the signaling network by literature research on interactions. This reconstruction resulted in a transcriptional regulatory network (TRN) which included two parts: the extracellular component (cytokines, chemokines and metabolic byproduct present in a tumor microenvironment) and the intracellular component (the master transcription factors (TF) implicated in macrophage polarization). We simulated the dynamics of our TRN using a Boolean approach. Then, Boolean attractors obtained from this network were classified into two main groups: pure (experimentally proven macrophages) or hybrid states (hypothesized phenotype found using boolean models and artificial intelligence [2, 3, 30]. In global terms, we used the following basis to label the attractors found in our simulation. For the M1 macrophages, we considered that the TFs associated with these phenotypes were STAT1 and NFκB [13, 18]. The activation of at least one of these TFs is implicated with the secretion of cytokines that eliminate the possibility of tumor cells to thrive and enhance a pro-inflammatory condition [23, 29]. M2 activation has been shown to be associated with pro-tumorigenic outcomes. The key TF implicated in M2a Th2 response and profibrotic is STAT6 [14, 27]. STAT6, at the same time, can inactivate the functions of NFκB and STAT1 through SOCS1, avoiding the M1 phenotype. The activation of the M2b macrophage is quite complex, with the inclusion of immune complexes (like immunoglobulin G (IgG)) and interleukin-1 receptors (IL1-R) (for example IL1-β) [4, 38]. The M2b macrophage has the ability to secrete not only anti-inflammatory cytokines but pro-inflammatory cytokines [28, 22]. The M2c macrophage emerges from the stimulation of IL-10 (the most important cytokine with an anti-inflammatory capacity and associated with bad prognosis in cancer [36, 43]). STAT3 is the condition for the anti-inflammatory capacity and inhibitor of the secretion of pro-inflammatory cytokines [12]. Finally, the M2d macrophage is activated via a co-stimulation of adenosines and TLR4, as well as the activation of HIF1-α [9, 1]. This macrophage is implicated in processes like angiogenesis and tumor progression, which is why they are also known as tumor-associated macrophages [9, 6]. The full description and characterization of the network used in this work is in [2]. The list of Boolean attractors and their classification in macrophage states can be reviewed in the supplementary material. The epigenetic landscape was built to apply a method that reduces the number of variables to a bi-dimensional map, in this case we used the distributed stochastic neighbor embedding algorithm (t-SNE). For the depth of the valley of each of the 13 phenotypes we used the value of the basin of attraction.

### 2.2. Stochastic epigenetic model

Our model is a stochastic partial differential equation (PDE) with gradient flow structure for the epigenetic potential (landscape) described above in an appropriate infinite-dimensional context and driven by an additive Wiener (Brownian) process. We consider perturbations of the form *σB*, where *B* is a Brownian process (in infinite dimensions) and *σ >* 0 is a constant. In what follows, we will refer to *σ* as the noise magnitude. We study all the necessary rigorous mathematical results for this stochastic system in [24], relying heavily on the infinite-dimensional results of stochastic analysis, ergodic theory, large deviations, and stochastic optimal control. We also present the details of our stochastic numerical methods in [24].

We remark on the significance of the epigenetic potential (shown in Fig. 1) in our study: it encapsulates the relevant biological information for our system. Hence, we have stored the data and code necessary to obtain such potential in a repository (see Data Availability Statement) along with the Python code for the evolution of the stochastic epigenetic system.

**Figure 1:**
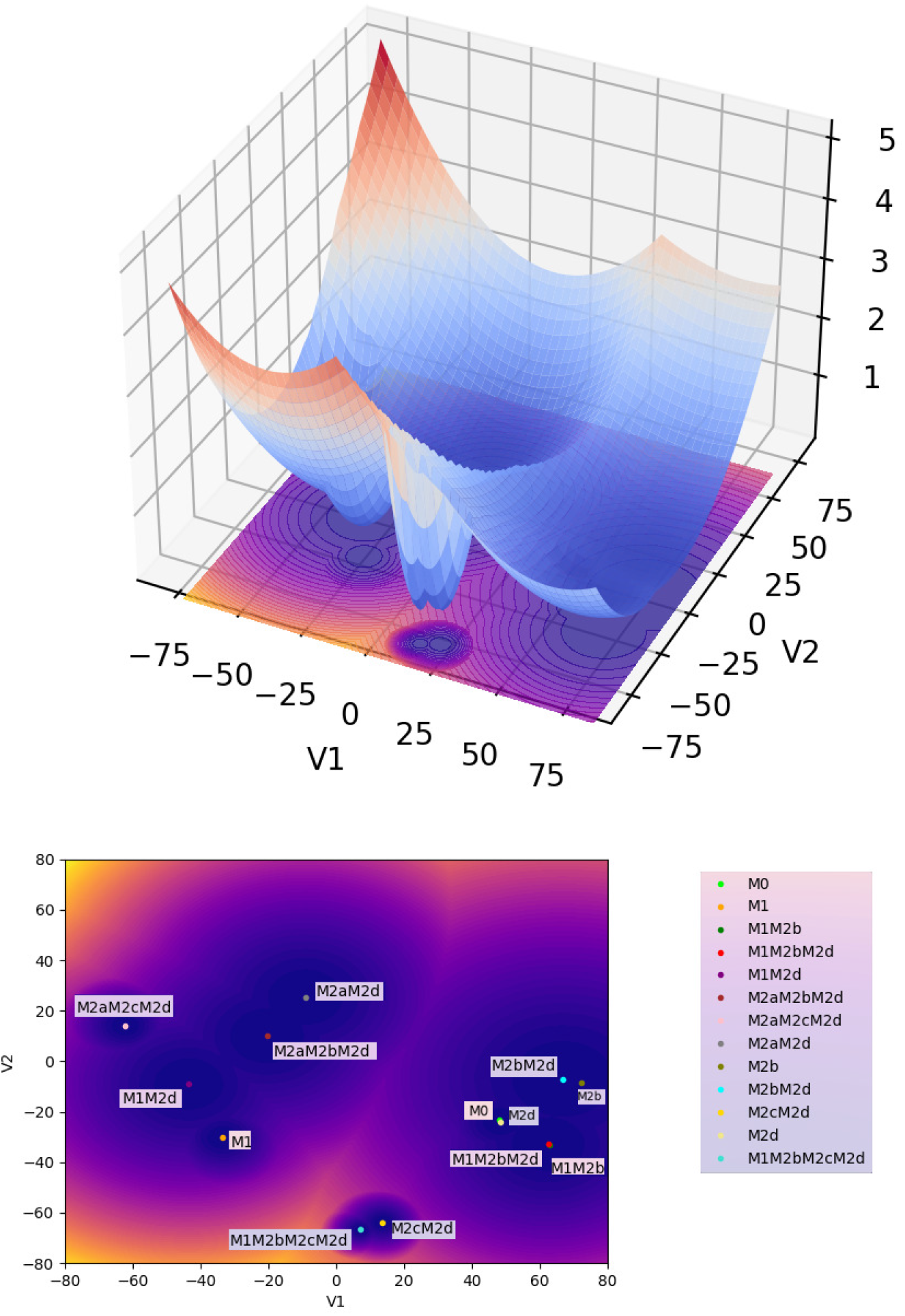
Macrophages’ epigenetic potential (Waddington’s landscape) in 3D (top) and the corresponding contour representation (bottom). The figure at the bottom shows the locations of the thirteen types of macrophages (phenotypes) in the epigenetic landscape, where each dot is the center of a phenotype, and the contours represent the depth of each phenotype (dark blue corresponds to the deepest region). In what follows, we use the contour representation of the epigenetic potential.

## 3. Results

We compared the behavior of all types of macrophages immersed in different values of (the magnitude of the) noise (*σ*); this noise represents the environmental fluctuations in a TME. Lower noise values in our study will represent a starting TME, corresponding to a nascent tumor still hijacking the processes in its surroundings. Meanwhile, as the noise increases, the microenvironment evolves to a much more complicated scenario, where the tumor is already thriving with the surroundings hijacked into a tumor-supportive microenvironment. Fig. 1 depicts the locations of the thirteen types of macrophages (phenotypes) in the epigenetic landscape, where each dot is the center of a phenotype, and the contours represent the depth of each phenotype (dark blue corresponds to the deepest region). These two-dimensional projections of each phenotype and the associated contours represent attractors determined using a boolean approach from [2]. We can observe 5 clusters, where 4 have macrophages with a tumor-supportive function, and only one cluster has a tumor suppressor function (M1 and M1M2d).

In the following simulations, we used different starting points in the landscape of the TAMs to evaluate how the noise could affect the dynamics (epigenetic state) of the adaptation of the macrophages to distinct stages of a tumor microenvironment. We ran our simulations with different noise magnitudes but for the same time interval. The first scenario starts with an M1 macrophage phenotype associated with the capacity to eliminate tumor cells. Fig. 2 shows that the macrophage remains in the basin of attraction of M1 for the entire time interval of the simulations. It is significant to notice the effect of the magnitude of the noise in the distribution of the epigenetic state of the macrophage: as the magnitude increases, the region within the M1 basin corresponding to the state of the macrophage (white area) gets more spread out. We proved in [24] that this distribution converges to a location concentrated at the basin’s center (yellow dot) as the noise decreases, showing the expected behavior in a noiseless controlled environment. The situation changes drastically in Fig. 3: for this noise magnitude (a more advanced stage of the TME), the macrophage remains for a short time in the M1 basin until it suddenly jumps to the M1M2d basin and, as time progresses, the macrophage keeps moving through the epigenetic landscape until reaching the basing corresponding to the M2aM2d phenotype. We call these transitions (entirely driven by the TME) noise-induced polarization. Although our model is stochastic with a driving random component (the TME), we can estimate, in a rigorous mathematical setting with probability 1, the exit time and the transition points in Fig. 3. This is relevant since these transitions can only occur in the so-called mountain passes of the epigenetic landscape, showing that even though the environmental epigenetic fluctuations drive the system, also the genetic component (encapsulated in the epigenetic landscape) determines the time and region of the transitions (polarizations, epigenetic mutations). We also remark that the results in Fig 2 do not indicate that, for lower noise magnitudes, there will not be a transition (polarization). We can prove, with probability 1, that such transitions will occur. However, it will be complicated to observe them computationally: the transition time becomes larger since it depends inversely on the noise magnitude. These results may also show the possible relevance of our methodology in the study of aging [37] and diseases like Alzheimer’s [31].

**Figure 2:**
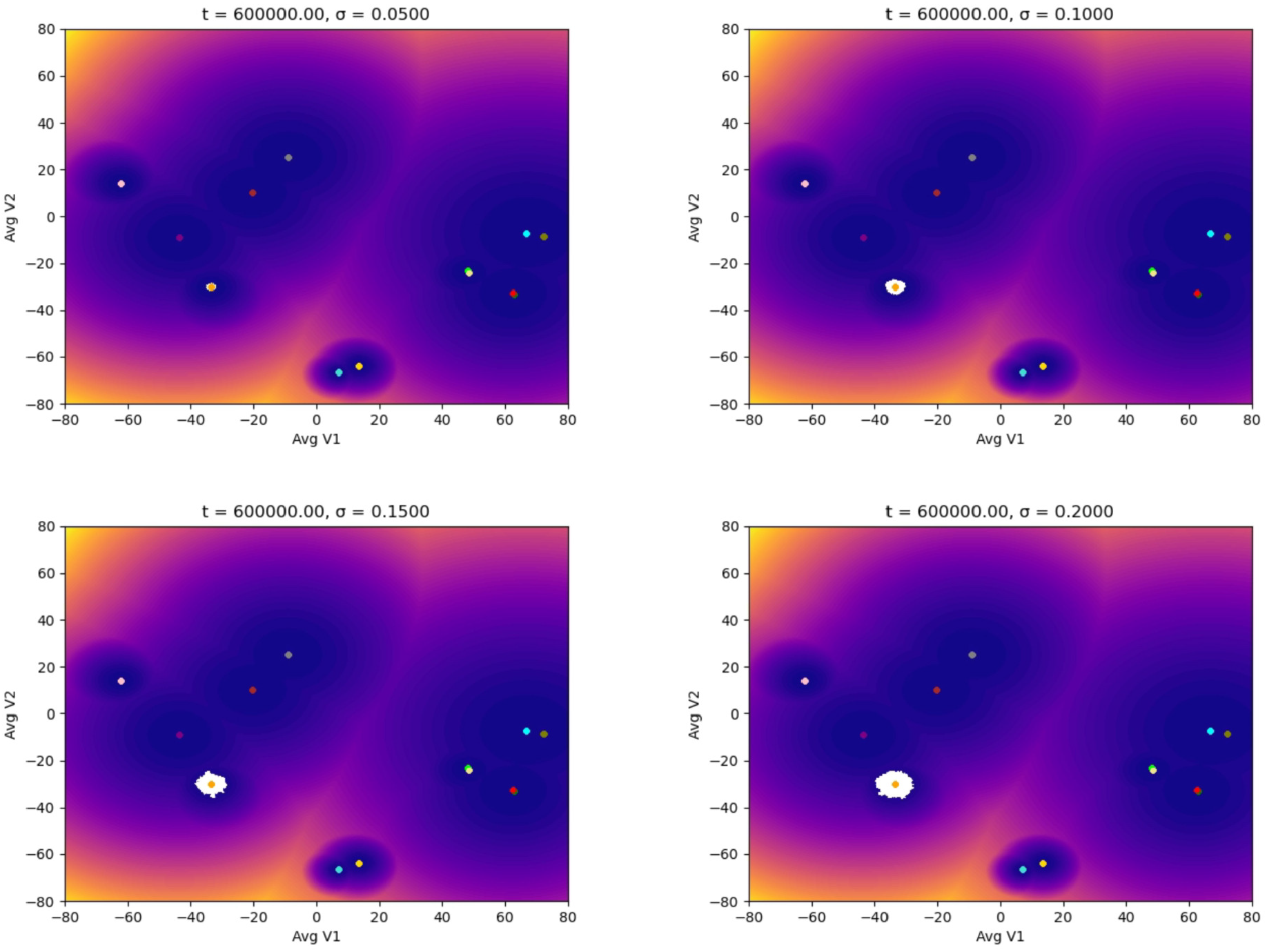
Evolution of the epigenetic state of a macrophage (in white) starting at the M1 phenotype center, up to *t* units of time, and subjected to different noise magnitudes, *σ*, representing the stages of the TME: the higher the value of *σ*, the more advanced the tumor is. The noise (a Brownian process) represents the molecular signals in the TME.

**Figure 3:**
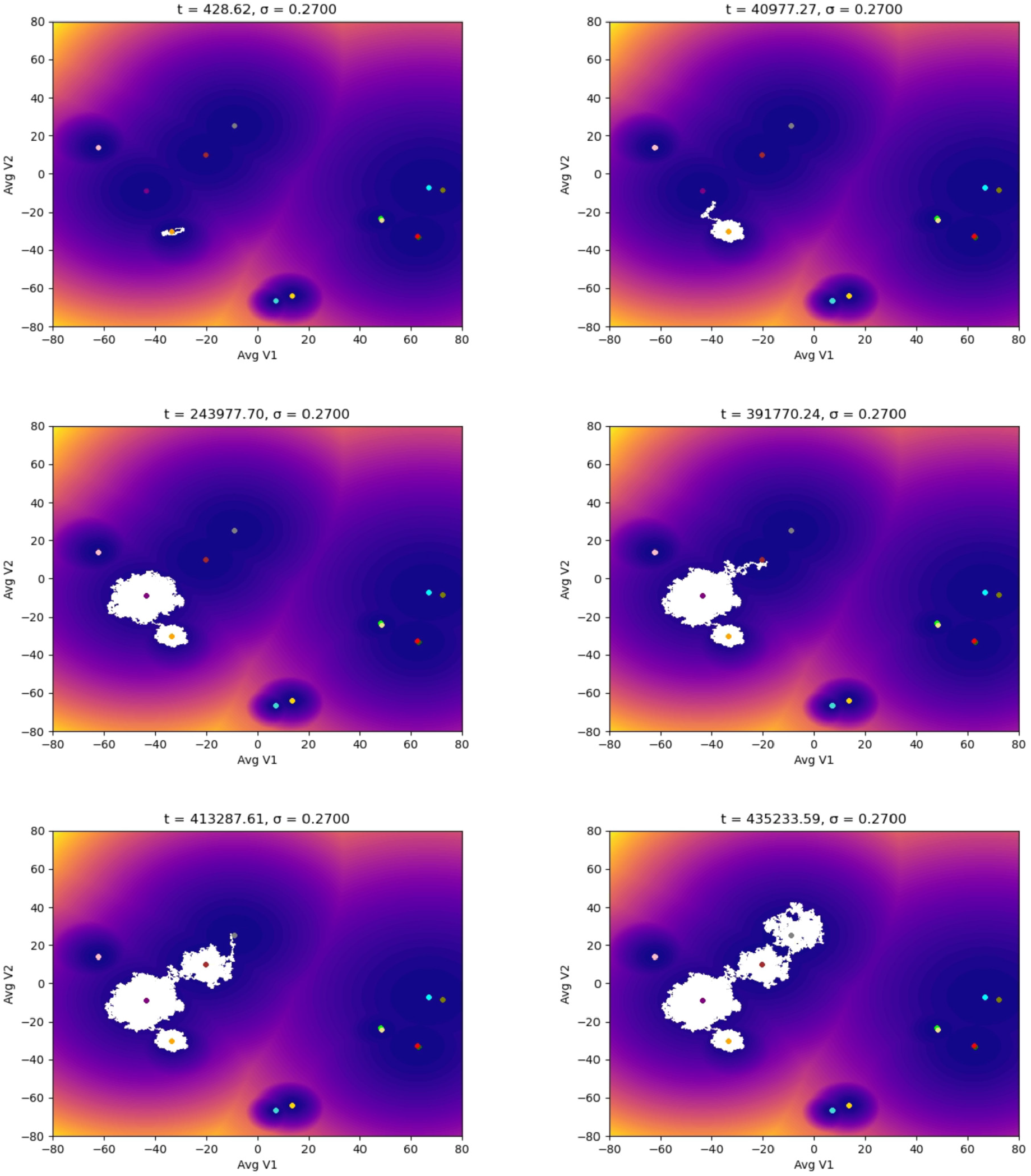
Evolution of the epigenetic state of a macrophage (in white) starting at the M1 phenotype center showing the noise-induced polarization into protumoral phenotypes due to the TME. Each plot shows the system’s dynamics up to *t* units of time, with fixed *σ* = 0.27.

Despite having the compartment M2d implicated in the secretion of components associated with tumor promotion, M1M2d has a role in wound healing [41, 5]. Wound healing is a significant counterbalance provided by the secretion of inflammatory components from the M1 phenotype. As we mentioned before, Fig. 3 shows that the M1M2d macrophage finally polarizes into the cluster that holds two tumor-supportive phenotypes (M2aM2d and M2aM2bM2d); in this scenario, the macrophage will only secrete components that will benefit a microenvironment that will allow the tumor to grow.

Macrophages enter the location where the tumor resides as monocytes circulating in the blood. Figs. 4 and 5 show the simulations where we were interested in studying how the monocyte responds to the perturbation from the TME by increasing the noise. We can observe that the monocyte polarizes into an M2d macrophage despite a low noise value. As the noise and the time increase, the monocyte polarizes into a macrophage with an M2 phenotype implicated in a tumor-supportive role, finalizing in the M2bM2d phenotype, which favors tumor progression, metastasis and immune evasion [2, 3]. Finally, Figs. 6 and 7 show our results starting from the hybrid phenotype M1M2bM2cM2d.

**Figure 4:**
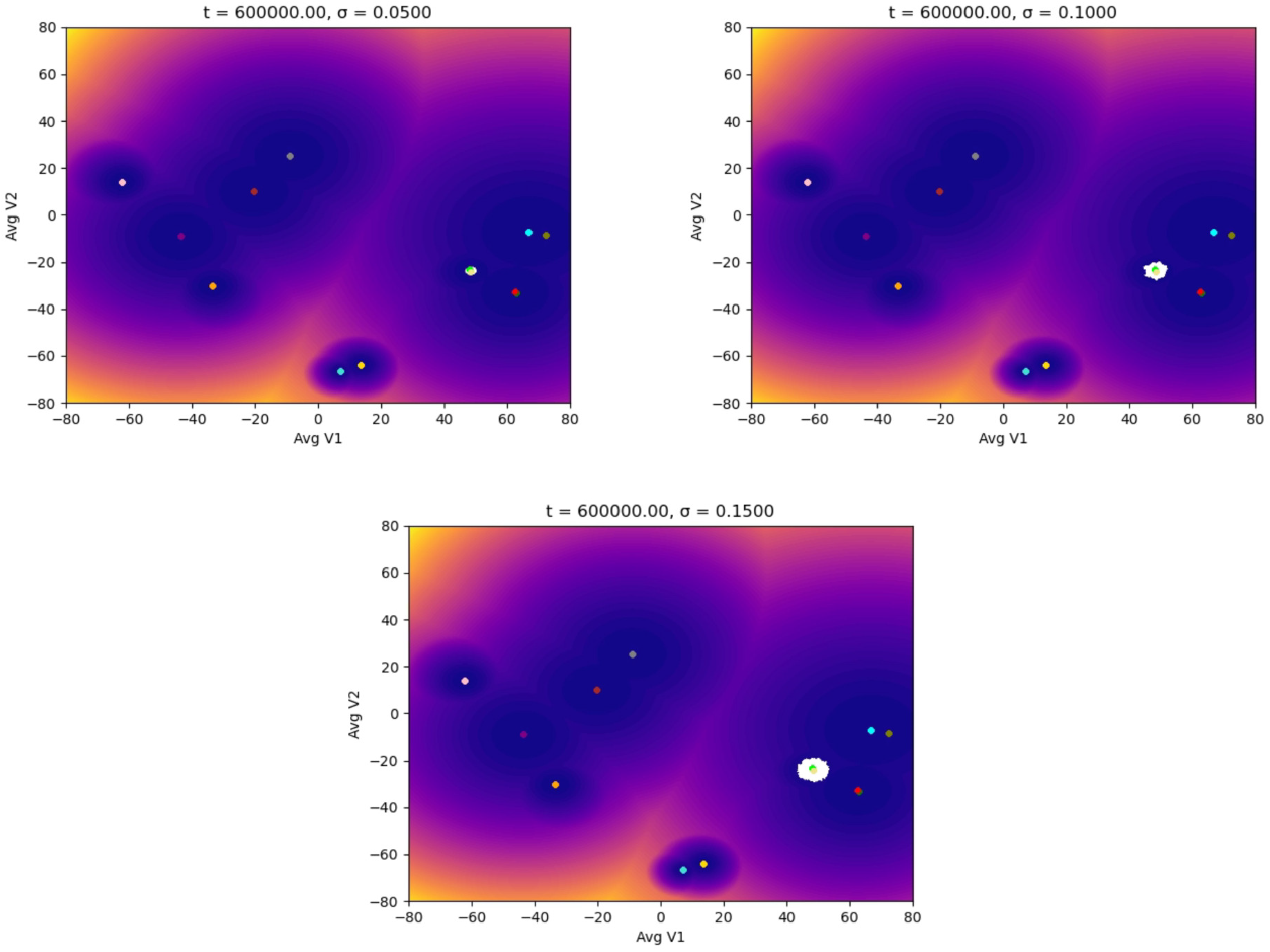
Evolution of the epigenetic state of a macrophage (in white) starting at the M0 phenotype center (monocyte), up to *t* units of time, and subjected to different noise magnitudes, *σ*, representing the stages of the TME: the higher the value of *σ*, the more advanced the tumor is. The noise (a Brownian process) represents the molecular signals in the TME.

**Figure 5:**
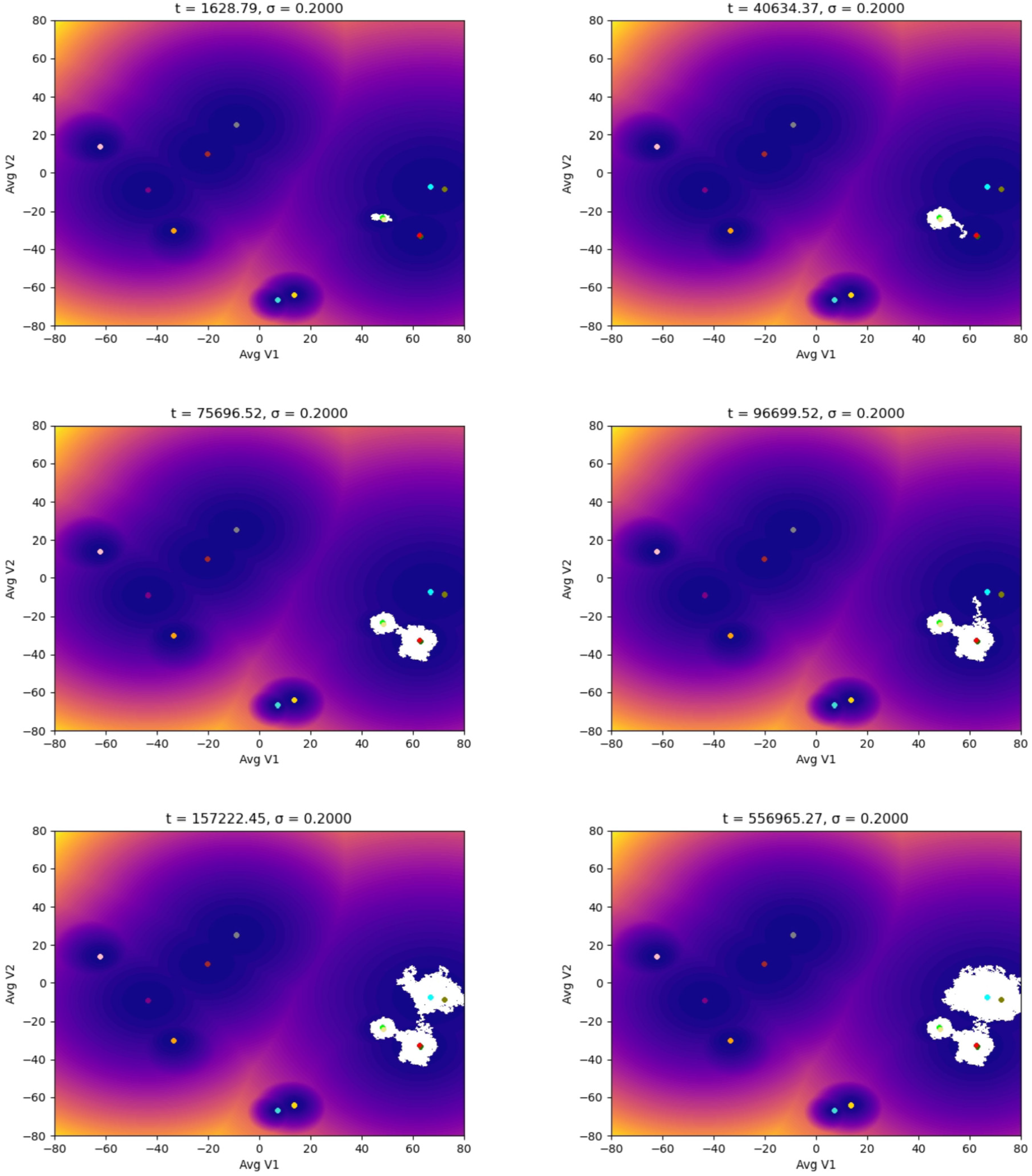
Evolution of the epigenetic state of a macrophage (in white) starting at the M0 phenotype center showing the noise-induced polarization into protumoral phenotypes due to the TME. Each plot shows the system’s dynamics up to *t* units of time, with fixed *σ* = 0.20.

**Figure 6:**
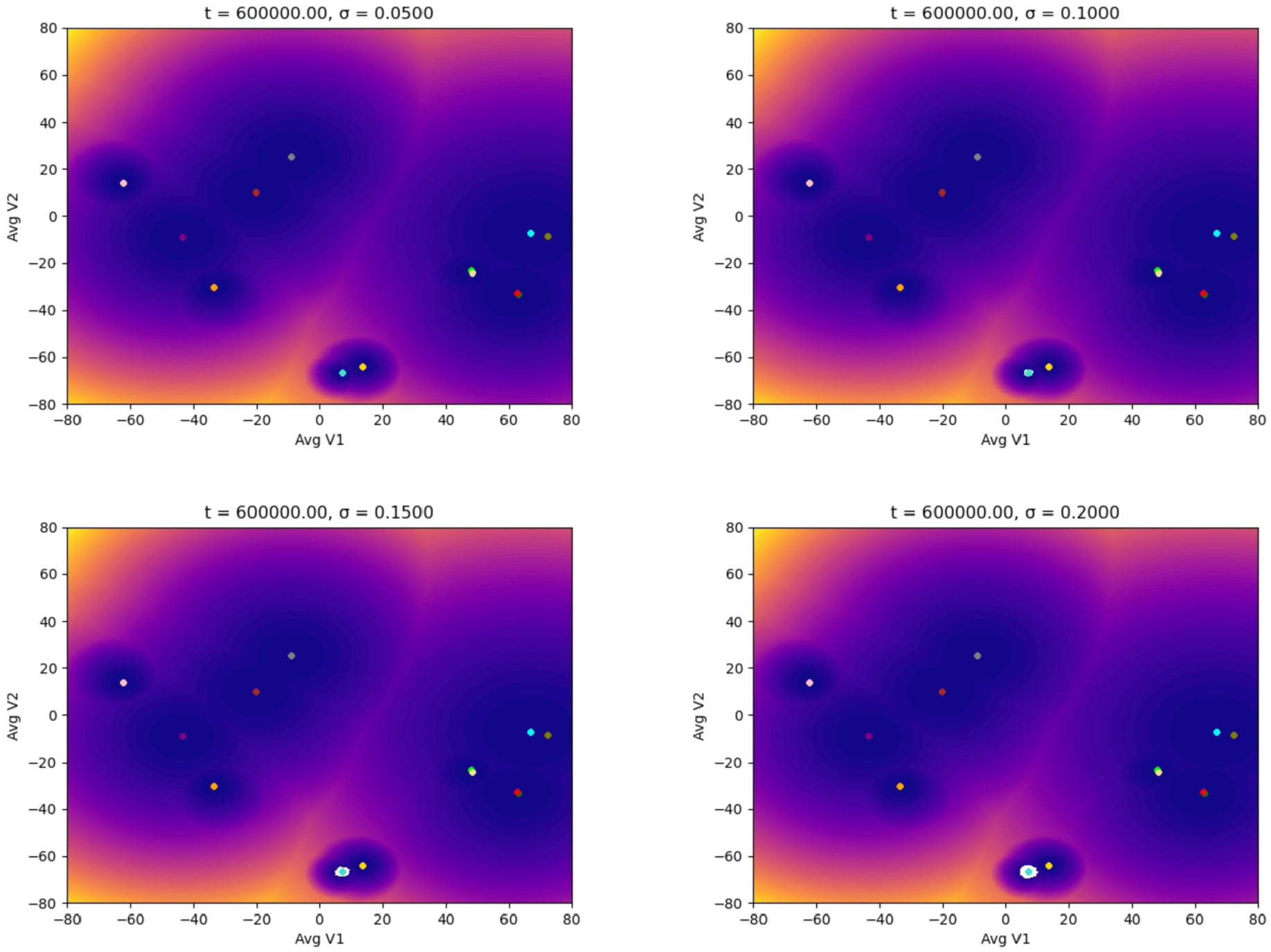
Evolution of the epigenetic state of a macrophage (in white) starting at the M1M2bM2cM2d phenotype center, up to *t* units of time, and subjected to different noise magnitudes, *σ*, representing the stages of the TME: the higher the value of *σ*, the more advanced the tumor is. The noise (a Brownian process) represents the molecular signals in the TME.

**Figure 7:**
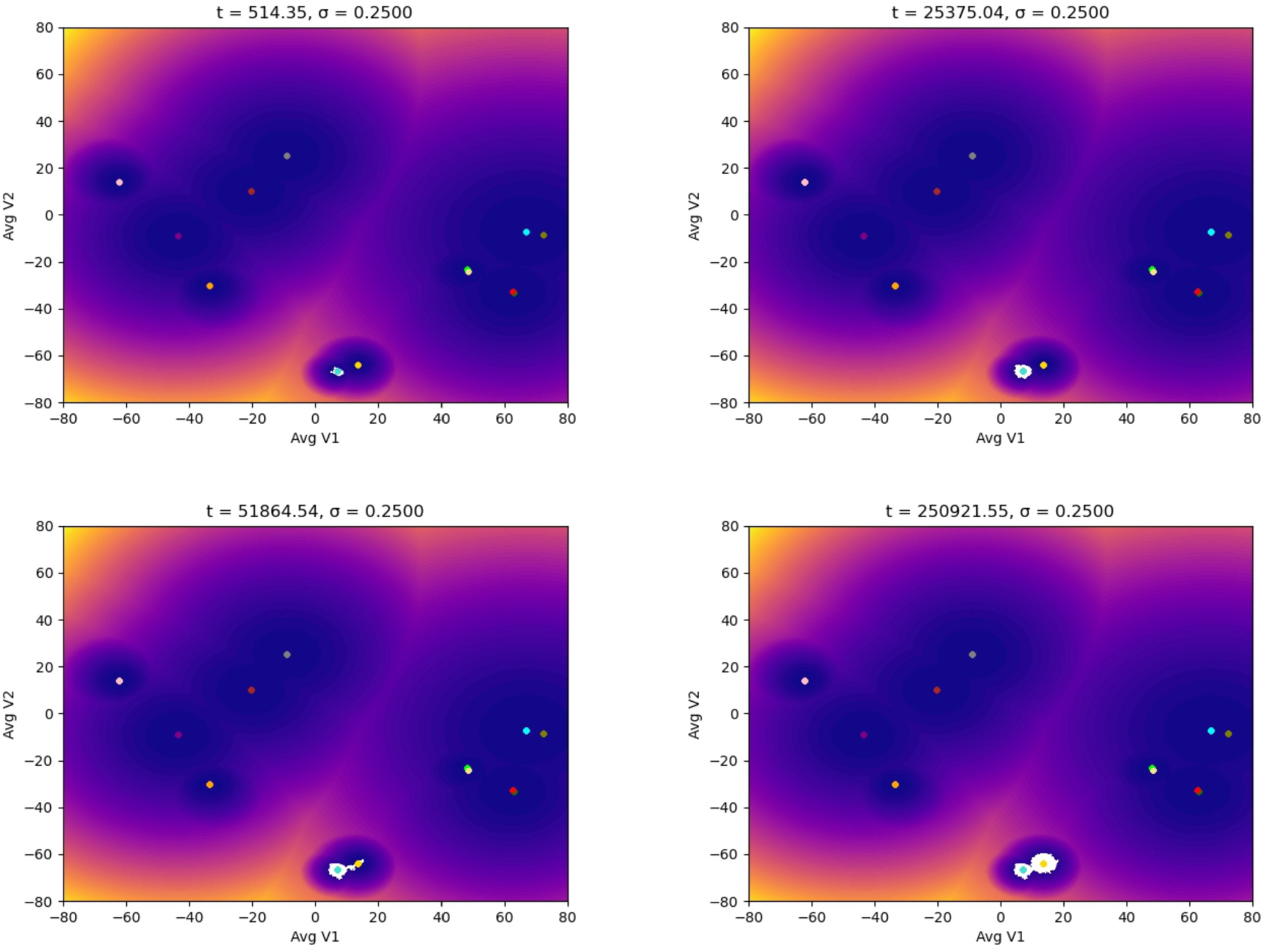
Evolution of the epigenetic state of a macrophage (in white) starting at the M1M2bM2cM2d phenotype center showing the noise-induced polarization. Each plot shows the system’s dynamics up to *t* units of time, with fixed *σ* = 0.25.

Our results show that macrophages can be associated with a poor prognosis in many types of cancer. Therefore, a therapy involving macrophages has to consider this situation and requires an additional mechanism to ensure the appropriate epigenetic stability; we will discuss this situation in detail in the next section.

## 4. Discussion

Our model shows that the influence of the TME on a therapy involving macrophages can render it ineffective and even benefit the tumor progression. Since the TME produces a noise-induced polarization of macrophages, we need an additional mechanism to promote epigenetic stability. [17] shows the unique role of p53 in the regulation of M2 macrophage’s polarization, in particular, a function for the p53/MDM2/c-MYC axis as a physiological ‘brake’ to the M2 polarization process; p53 is the first transcription factor reported to suppress M2 macrophage’s polarization. On the other hand, [10] demonstrates that increased p53 expression in TAMs induces canonical p53-associated functions such as senescence and activation of a p53-dependent senescence-associated secretory phenotype. This was linked with decreased expression of proteins associated with M2 polarization by TAMs. [10] also presents a clinical trial in patients with solid tumors combining APR-246 (which increases p53 signaling) with pembrolizumab, showing that biospecimens from select patients had suppression of M2-polarized myeloid cells and an increase in T-cell proliferation, leading to suggest that increasing p53 expression in TAMs ‘reprograms’ the TME to augment the response to immune checkpoint blockade.

The previous observations, along with our results about noise-induced polarization, highly suggest that any immunotherapy involving macrophages should include a mechanism capable of increasing p53 signaling, like APR-246 (eprenetapopt), to promote the appropriate epigenetic stability; see Fig. 8. We also mention the review by Levine and Berger [16] that discusses the cross-talk between p53 and epigenetic programs: p53 not only enforces the stability of the genome by preventing genetic alterations in cells but also plays a significant role in regulating the epigenetic changes that occur in cells.

**Figure 8:**
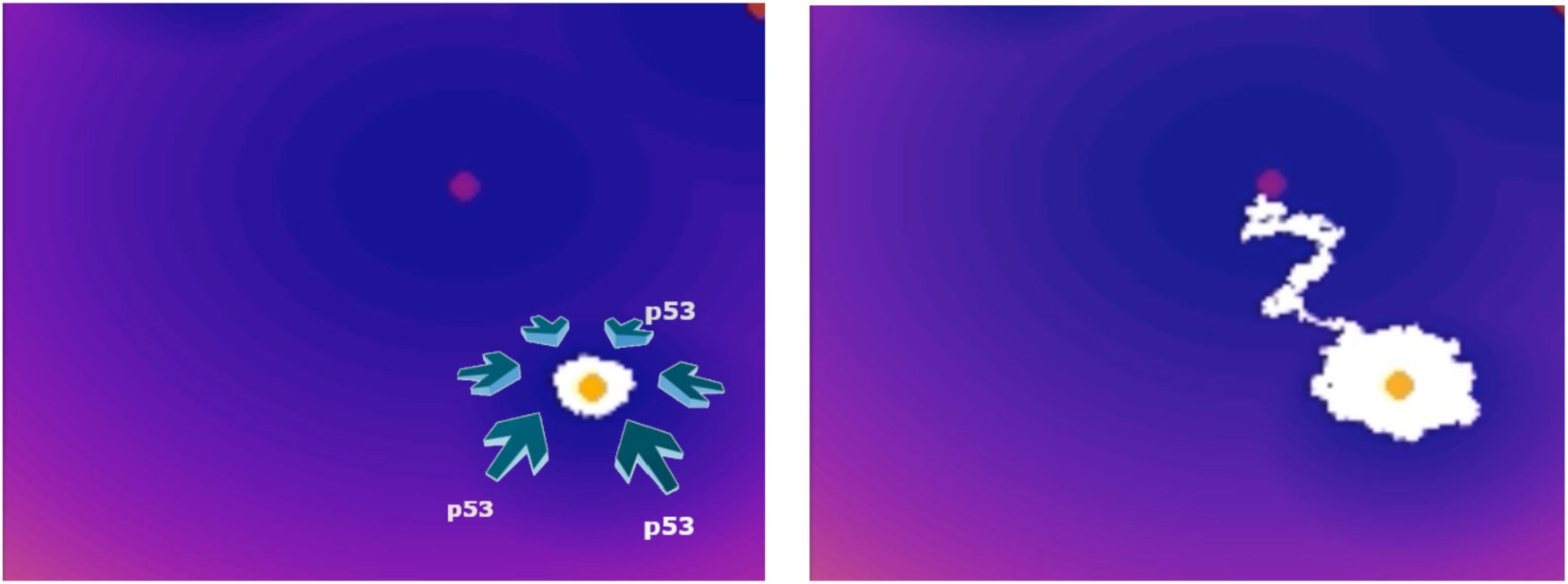
The effect of p53 on the epigenetic evolution of macrophages in a TME. Left: p53 stabilizes the epigenetic M1 state. Right: in the absence of an epigenetic stabilizer, the macrophage polarizes into a protumoral state.

Only targeting macrophage polarization as a treatment against cancer may not be enough, mainly because of how susceptible they are to the TME. Pembrolizumab, an immune checkpoint inhibitor targeting the PD-1 receptor, has shown considerable efficacy in activating T cells within the TME. Although its primary mechanism is aimed at T cells, emerging evidence suggests it may also affect macrophage polarization and function, which are crucial factors in the immune response to cancer. For example, in [11], the authors demonstrate that macrophages can express PD-1, particularly in the TME, where macrophages exhibit an exhausted phenotype, leading to decreased phagocytic activity and altered cytokine production. Inhibition of PD-1 on macrophages could rejuvenate their phagocytic abilities and shift their polarization towards a more pro-inflammatory phenotype. Because of the action of this drug on T cells, it could lead to the production of interferon-gamma, a cytokine that can induce M1 polarization of macrophages, supporting a more robust anti-tumor response [34, 8]. Furthermore, blocking PD-1 might alter the cytokine profile within the TME, reducing immunosuppressive cytokines (IL-10) that favor M2 polarization and increasing the cytokines that promote M1 polarization [21, 39].

On the other hand, APR-246 is a reactivator of mutant p53 protein that induces apoptosis and inhibits tumor growth in tumor cells; this restored p53 function can influence the TME, potentially altering the secretion of cytokines and chemokines that regulate macrophage behavior [15]. Moreover, APR-246 could indirectly affect macrophages by modulating the expression of genes involved in the immune response. For instance, p53 can influence the expression of various cytokines and surface molecules that may affect macrophage recruitment and polarization [10, 42]. Finally, APR-246 might promote a pro-inflammatory environment by enhancing p53’s role in apoptosis and cell-cycle arrest in tumor cells; this could lead to increased tumor cell death, releasing tumor antigens and associated danger signals that could shift macrophage polarization towards a pro-inflammatory, anti-tumoral M1 phenotype [33].

The combination therapy between pembrolizumab, APR-246, and macrophages opens up new avenues for cancer therapy, particularly in understanding how immune checkpoint inhibitors can modify the tumor microenvironment to promote more robust anti-tumor responses. A current clinical trial evaluates the combination of both drugs and how they affect the TME and the immune checkpoint inhibitors [33, 26]. The potential reprogramming of macrophages could amplify the efficacy of pembrolizumab and possibly improve outcomes in cancers where macrophages play a significant role in tumor progression and immune evasion. Further research is necessary to delineate how pembrolizumab and similar drugs affect macrophage polar-ization in different types of cancers in conjunction with other treatments. These findings could lead to new combination therapies that target multiple cells and pathways within the immune system for a more comprehensive approach to cancer immunotherapy. An appropriate understanding of these mechanisms requires more focused studies, which could lead to more effective strategies in immuno-oncology, enhancing the therapeutic benefits of drugs like pembrolizumab in various cancer types.

Our stochastic optimal control model in [24] shows that the effect of p53 can be considered as a control on the drift of the system. Moreover, as long as the macrophages stay in an antitumoral state, the noise in the TME should be reduced as a consequence of the macrophages attacking the cancer cells, which will also help with the epigenetic stability (see Fig. 2); we can study this phenomenon with our model through a control in the diffusion (noise) term, which can change with the state of macrophages, capturing the impact of macrophages on the tumor as they polarize from M1 to M2 states. On the other hand, M1 macrophages augment the response to immune checkpoint blockade and increase T-cell proliferation: since the interaction of these different agents is relevant to describe the tumor, we believe that our model can capture the complexity of the TME more accurately by considering, in addition, the theory of (stochastic evolutionary) games with predator-prey structure including many species (macrophages, T-cells, cancer cells, etc). In this case, we could study the population of the different species, their interactions, their epigenetic states, the influence of hypoxia, toxicity, nutrients, external controls (like APR-246, pembrolizumab, epidrugs), and the effect these factors have on the TME: notice that in this ecosystem the species may mutate and change their roles (phenotypes) drastically in a “short-term”; cf. evolutionary game theory [35], evolutionary invasion analysis (adaptive dynamics) [7]. We believe these aspects are fundamental in determining an appropriate immunotherapy for cancer.

## Supporting information

See Fig 3

See Fig 5

See Fig 4

See Fig 2

## Author contributions

The three authors contributed to initiating the study and overseeing the research activities. Jesus Sierra was involved in the formal analysis, investigation, methodology, software, and validation. Ugo Avila was involved in conceptualization, investigation, and verification. Pablo Padilla contributed to the formal analysis, methodology, validation, and project administration. The three authors contributed to the manuscript’s writing, review, and editing.

## Data availability statement

We have archived our Python code on Zenodo (https://zenodo.org/records/13314201). We perform extensive stochastic and numerical analysis of our epigenetic model in [24]. Data related to the epigenetic potential can be found at (https://github.com/resendislab/M1-M2-Macrophage-Polarization).

